# Acoustic and vibrational signaling in true katydid *Nesoecia nigrispina*: three means of sound production in the one species

**DOI:** 10.1101/2021.12.30.474575

**Authors:** Olga S. Korsunovskaya, Rustem D. Zhantiev

**Author notes:** Corresponding Author: Olga Korsunovskaya, *Leninskie Gory, 1, building 12, Moscow, 119234, Russia.

## Abstract

The males of Mexican katydids *Nesoecia nigrispina* (Stal, 1873) produce calling songs and protest sounds using the same stridulatory apparatus as in most of the other Ensifera at the base of the elytra. It includes pars stridens on the upper elytron and plectrum on the lower. Calling sounds of two types (fast and slow) are 2-pulse series, repeated with a frequency of 1.2-4.5 s^−1^. Protest signals in the form of short trills from the same pulse duration males produce with tactile stimulation. The pulse repetition rate is more higher than that of the calling sounds - up to 10 (mean c. 8) s^−1^. The frequency spectra of these signals have maxima in the band of 14–15 kHz. However, in addition to the sounds described, both males and females are capable to produce protest signals of the second type with the help of another sound apparatus, namely with the help of the wings. Insects with removed elytra are unable to produce an audible sound. Thus, the sound is produced by the friction of the wings on the elytra, but there are no specialized stridulatory structures on them. In individuals of both sexes, in response mainly to tactile stimulation, short clicks are recorded, which they make, apparently, by the mandibles. Vibrational signals at tremulation are emitted by individuals of both sexes during courtship and males, completing the calling signal cycle and after copulation. It is possible that vibrational signals are an additional factor in reproductive isolation in sympatric species, since the calling sound signals in representatives of the genus *Nesoecia* are similar and exhibit significant variability.

## Introduction

Katydids of a large subfamily Pseudophyllinae (true katydids) are common in the tropics and subtropics in both the Old and New Worlds. Many of them mimic the leaves of plants, but a large group of these insects has a cryptic coloration and they don't look like plants. The latter include representatives of the genus *Nesoecia*. They are large, earthy colored katydids with shortened wings. Currently, 8 slightly different morphologically species have been described that live in southern Mexico, Brazil, and one of the Galapagos Islands (Floreana) (Cigliano et al., 2021). Little is known about the biology of true katydids. In some species of the tribes Cocconotini, Pleminiini and some others acoustic signals have been studied, which in some cases are supplemented or accompanied by vibrations produced by tremulation or substrate drumming (Belwood, Morris, 1987; Belwood, 1990, Morris et al., 1994, Roemer et al., 2010; Rajaraman et al., 2015). Some species possess unique sound apparatus, such as the females of *Panopnoscelis specularis* Beier (Montealegre-Z et al., 2009) or sound emission mechanisms unusual for katydids, such as the abdomino-alary in *Pantecphylus cerambycinus* Karsch, 1891 (Heller, 1996). In this study we used specimens of *Nesoecia nigrispina* from a laboratory culture. This made it possible to study in detail their behavior and signaling using both sound and vibratory signals, and to identify in this species three means of eliciting sounds that insects use in different behavioral contexts.

## Materials & Methods

We used *Nesoecia nigrispina* (Stal, 1873) (Orthoptera, Pseudophyllinae, Cocconotini) from the laboratory culture of the Moscow Zoo and the Department of Entomology of Moscow Lomonosov State University. The basis of the culture was a female caught in Mexico, in the northern half of the Yucatan Peninsula (Chechen Itza), under the bark of a tree and transferred to the insectarium of the Moscow Zoo. The lifetime of one generation is about a year, but in nature its duration is apparently shorter, because in the last 1-1.5 months of life, insects become inactive, sedentary and often lose one or two legs, which, obviously, makes them easy prey for predators. One female, living for at least 6 months, mates and lays eggs for most of her life. The development of larvae at 25°C lasts about four months. By now, 5-6 generations have already passed - the descendants of the first female. Katydids were kept in cages measuring 30×30×30 cm. In one container, both adults and nymphs of different ages could be kept. Cannibalism is not typical for this species. The fodder was raspberry, blackberry, oak, lettuce leaves, in the summer - clover and dandelion leaves, slices of fruit (orange, apple), oatmeal and protein supplements in the form of dried freshwater amphipods of the genus *Gammarus*. For laying, females were offered containers with a wet mixture of peat and soil. *N. nigrispina* are nocturnal. We did not register during the daytime singing. During daylight hours, insects sit on the walls of cages or pieces of wood, as a rule, completely motionless. Katydids are reared at temperature of 25–27^о^С and under a constant photoperiod 12 L:12 D.

The sound signals of *N. nigrispina* were digitally recorded using a Roland R-05 recorder (frequency response 0.02–40 kHz, flat response 0.02–20 kHz) with a sampling rate of 96 kHz or using Brüel and Kjær microphone 4135 and amplifier 2606 (frequency response flat up to 40 and 70 kHz respectively) with a sampling rate of 200 kHz. Amplifier was connected via an analogue-to-digital converter E-14-440 (L-Card, Russia) to a PC. The vibratory signals of *N. nigrispina* were recorded using a ZPK-56 pickup head (Russia) with a piezoceramic element made of barium titanate (frequency response 30–12000 Hz) mounted horizontally with the needle stand upside down. This device was connected via the same analogue-to-digital converter to PC. Digitizing frequency at recordings of vibrational signals was 5 kHz. The signals were recorded using the program PowerGraph 3.3. (DiSoft, Russia). Acoustic and vibratory signals were processed using the CoolEditPro 2.1 and PowerGraph 3.3 softwares.

Night recordings were performed using a Nikon 1J4 video camera without an infrared filter under the light of an Orient SAL-130C infrared illuminator. High-speed video filming was carried out with a frame rate of 1000 per sec.

Scanning electron micrographs (SEMs) of specimens were made with a JEOL/EO scanning electron microscope (Japan) (Electron Microscopy Laboratory, Faculty of Biology, Moscow Lomonosov State University). Photographs and measurements of morphological characters were made with a using a Canon EOS 6D digital camera with a Canon MP-E65 macro lens.

Sound and vibrational signal recordings were performed at 25-26 °C. Data are given as mean ± standard deviation, n≥20 per series.

### Terminology

At description of sound and vibratory signals we mainly follow bioacoustic terminology used Heller et al., 2021: calling song – spontaneous song produced by an isolated male. Syllable – the sound produced by one complete up (opening) and down (closing) stroke of the forewings (tegmina). Hemisyllables parts of syllable corresponding to each uni-directional movement of tegmina. Tooth-impact – short sound impulse arising during contact of a single tooth of the stridulatory file (pars stridens) with the plectrum. Click – fast train of sound waves arising at strucking any structures. Series – group of several syllables.

## Results

Proceeding from the fact that morphologically the Mexican and South American species of *Nesoecia* are extremely close, we considered it necessary to describe some characters of the male and female katydids, which we identified as *N. nigrispina*.

### Male

(Fig. 1 A). Body color brown, sometimes olive brown. Head round, fastigium triangular with sinuate apex, frons with two dark spots in the upper third (Fig. 1 E, F).

**Figure 1.**
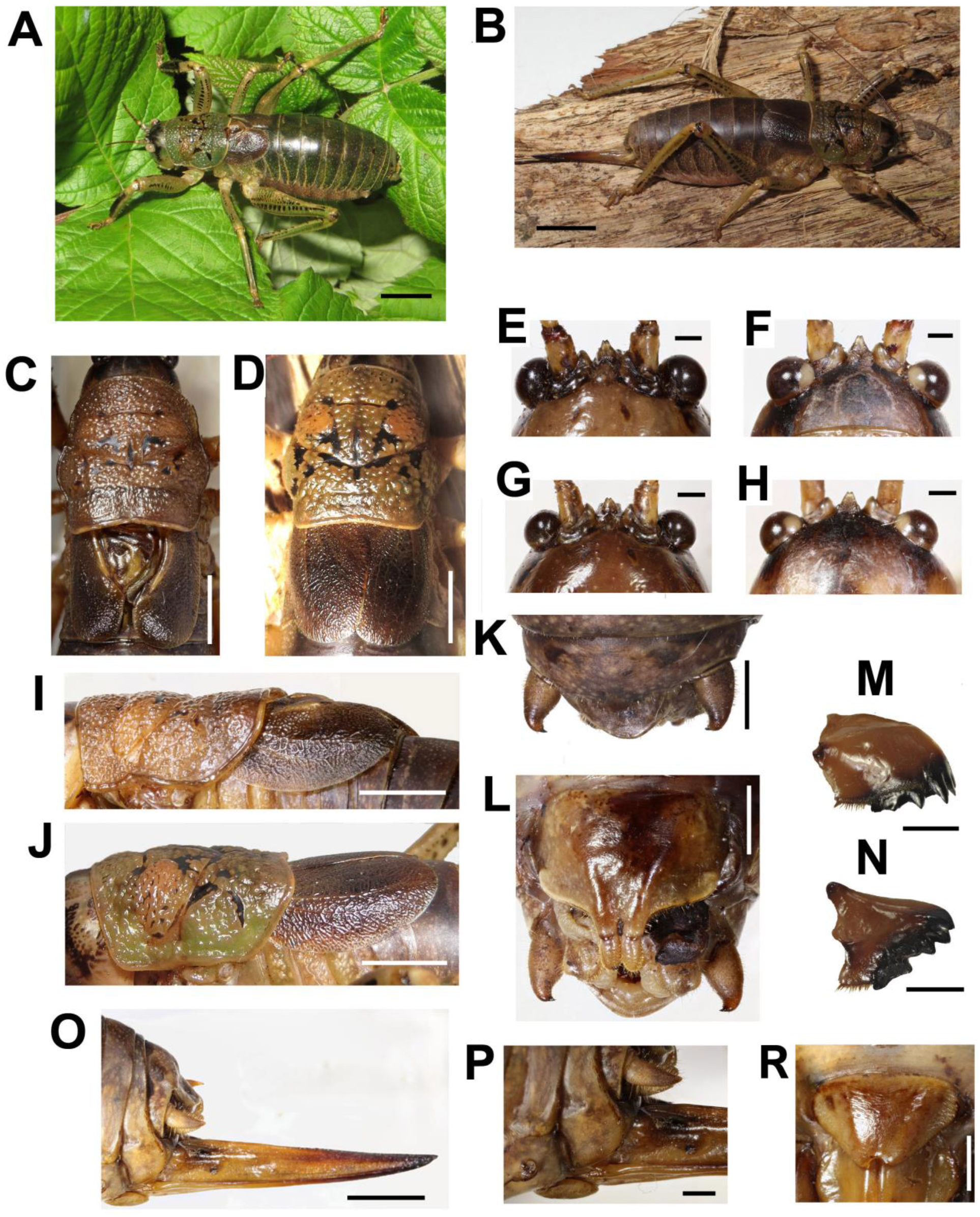
Habitus and body details of the male and female of *Nesoecia nigrispina*: A - male habitus, B - female habitus. C – male pronotum and tegmina, dorsal view; D – female pronotum and tegmina, dorsal view; E –male vertex and fastigium, anterior view; F – the same, dorsal view; G - female vertex and fastigium, anterior view; H – the same, dorsal view; I - male pronotum and tegmina, lateral view; J – female pronotum and tegmina, lateral view; K-male anal plate and cerci; L – male subgenital plate with stili; M – female left mandible, outside view; N – female right mandible, inside view; O – ovipositor; P - female’s terminalia: cerci and subgenital plate, lateral view, R – female subgenital plate. Scales: 10 mm (A, B), 5 mm (C, D, I, J, O), 2 mm (K, M, N, P, R), 1 mm (E-H)

Pronotum coarsely granulated with black spots and short interrupted bands (Fig. 1 C). Lateral lobes of pronotum wider than high (Fig. 1 I). Lower margin of lateral lobes with small notch before the middle (Fig. 1 I). Tegmina dark brown. Cell below the stridulatory file yellowish (Fig. 1 C, I). Wing apices reach the end of the 2^nd^ abdominal tergite. Legs with spines on lower surface. Lateral surfaces of femora and partly of tibiae with lines of dark spots. Anterior surface of fore tibia dark brown or black (Fig. 1 A). Anal plate wide-triangular, cerci curved inward at apex, their length approximately 1.5 times their maximum width, bear one short apical spine (Fig. 1 K). Subgenital plate is transverse; its width is 1.5 times its height; bifurcated distal part is stretched out and bears styli (Fig. 1 L).

Length, mm: body 41-47, pronotum length, 11-12, hind femora 15-17 (n=5).

### Female

(Fig. 1 B). Coloration and granulation as in males. Head, and frons as in males (Fig. 1 G, H). Pronotum is more olive comparing with males (Fig. 1 D, J). Lower margin of lateral lobes without notch before the middle (Fig. 1 D). Tegmina single color, brown, do not reach the end of the 2^nd^ abdominal tergite. Leg coloration as in males. Ovipositor straight (Fig. 1 O). Its length is approximately equal to that of the hind femur. Subgenital plate wide triangular, apically with small notch (Fig. 1 R).

Length, mm: body 47-52, pronotum length 11.5-13, hind femora (lower edge) 19-22, ovipositor (from subgenital plate to the tip) 18-20 (n=5).

#### Acoustic signals

*N. nigrispina* emit sound signals in three ways: using the tegminal sound apparatus (males only), wings, and, presumably, mandibles (individuals of both sexes).

Male calling songs are long sequences of series, consisting of two syllables. Insects produce them with the help of a stridulation apparatus, the structure of which is typical for the tegminal sound apparatus of katydids. On the upper tegmen, there is a stridulatory file (pars stridens) (Fig. 2 A, B), bearing 127 teeth rubbing against the plectrum at the medial edge of the lower elytron (Fig. 2 C). There is a resonating structure on it - a mirror, while the vein, which plays the role of pars stridens on the symmetrical tegmen, is thickened, but does not bear teeth (Fig. 2C).

**Figure 2.**
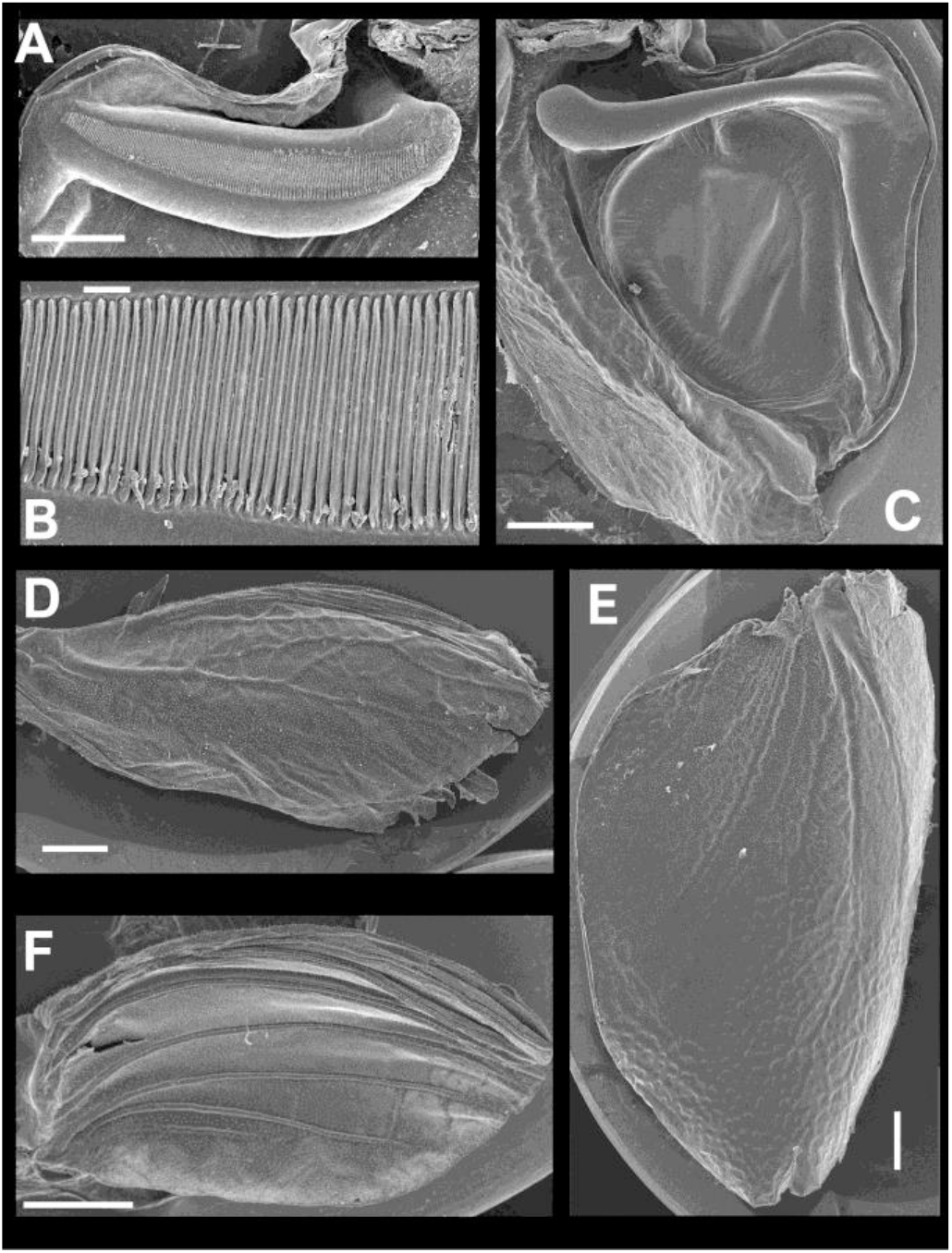
SEM images of the structures providing sound emission in *N. nigrispina*. (A, B) Male stridulatory file. (C) Male right tegmen, ventral view. (D) Right female wing, dorsal view. (E) Right female tegmen, ventral view. (F) Right male wing, dorsal view. Scale 1 mm (A, C-E) and 100 μm (B).

The calling of the male can continue for several hours. Males are capable to emit calling song with two rhythms: fast (Fig. 3 A, B) and slow (Fig. 3 C, D), and at the beginning of the signal with a greater duty cycle, the insect, as a rule, produces several series or several tens of series with a fast rhythm. With the simultaneous singing of several katydids, their acoustic interaction is observed, which is expressed in i) acceleration of the rhythm of signal emitting, ii) alternation, iii) synchronization), iv) emitting signals identical in amplitude-temporal pattern with tegminal protest sounds. As shown by high-speed video filming, the male produces a series of calling signal with two cycles of opening-closing of the tegmina, while after the initial tegminal opening, complete closure does not occur at the end of the first syllable. It is observed only after the second syllable. The form and duration of the syllable in the series, following in a fast and slow rhythm, is different. In fast rhythm series, the rise and fall times of the amplitude of the first and second syllables are approximately the same, the duration of the first syllable is 44.7+2.1 ms, the second one – 43.4+3.2 ms. The repetition rate of the series can fluctuate within 3–4.5 s^−1^, on average it is 4.2+0.2 s^−1^. The syllable repetition rate in the series averages 16.5+0.3 s^−1^. In the slow calling signal, the duration of the first syllable is 96.5+4.1 ms, the second one – 47.2+3.9 ms, and the rise time of the amplitude of the first syllable is much longer than the time of its fall and takes about 2/3 of the syllable duration. The shape of the second syllable is approximately the same as in a series with a fast rhythm. The repetition rate of the series can vary within 1.2–2 s^−1^, mean is 1.86+0.25 s^−1^, and the syllable repetition rate in the series is 9.09+0.28 s^−1^.

**Figure 3.**
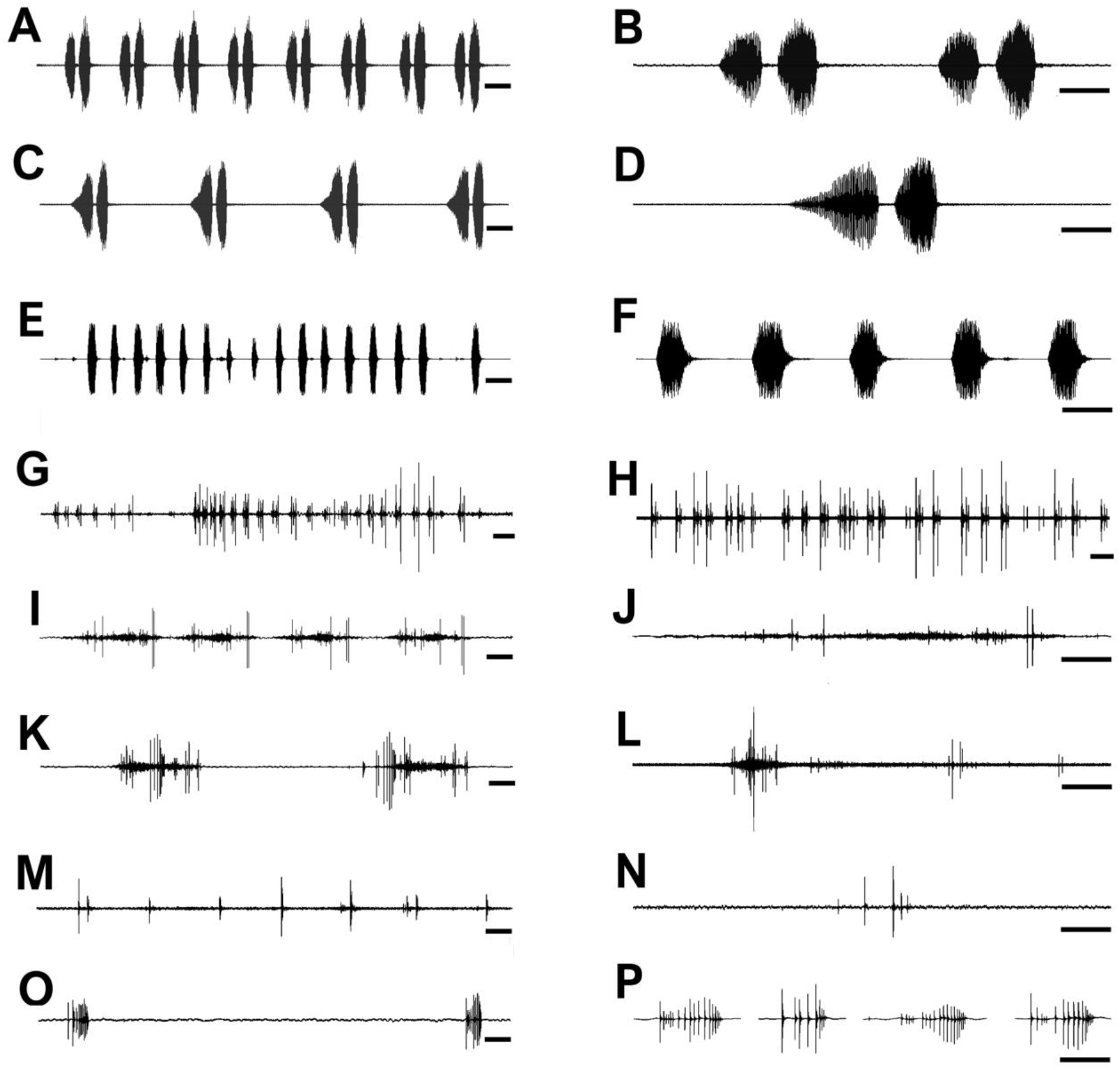
Oscillograms of the sound signals of *N. nigrispina*. (A, B) Fast calling signal. (C, D) Slow calling signal. (E, F) Tegminal protest sounds. (G, I, J) Male wing protest sounds. (H, K, L) Female wing protest sounds. (M, N) Male mandibular protest sounds. (O, P) Female mandibular protest sounds. Time scale: 50 ms (B, D, F, J, L, N, P), 100 ms (A, C, E, I, K, M, O), 1 s (G, H)

The frequency spectra of calling signals produced in fast and slow rhythms are similar: the main components lie in the range of 10-20 kHz, the dominant frequencies are c. 14-16 kHz (Fig. 4 A, B).

**Figure 4.**
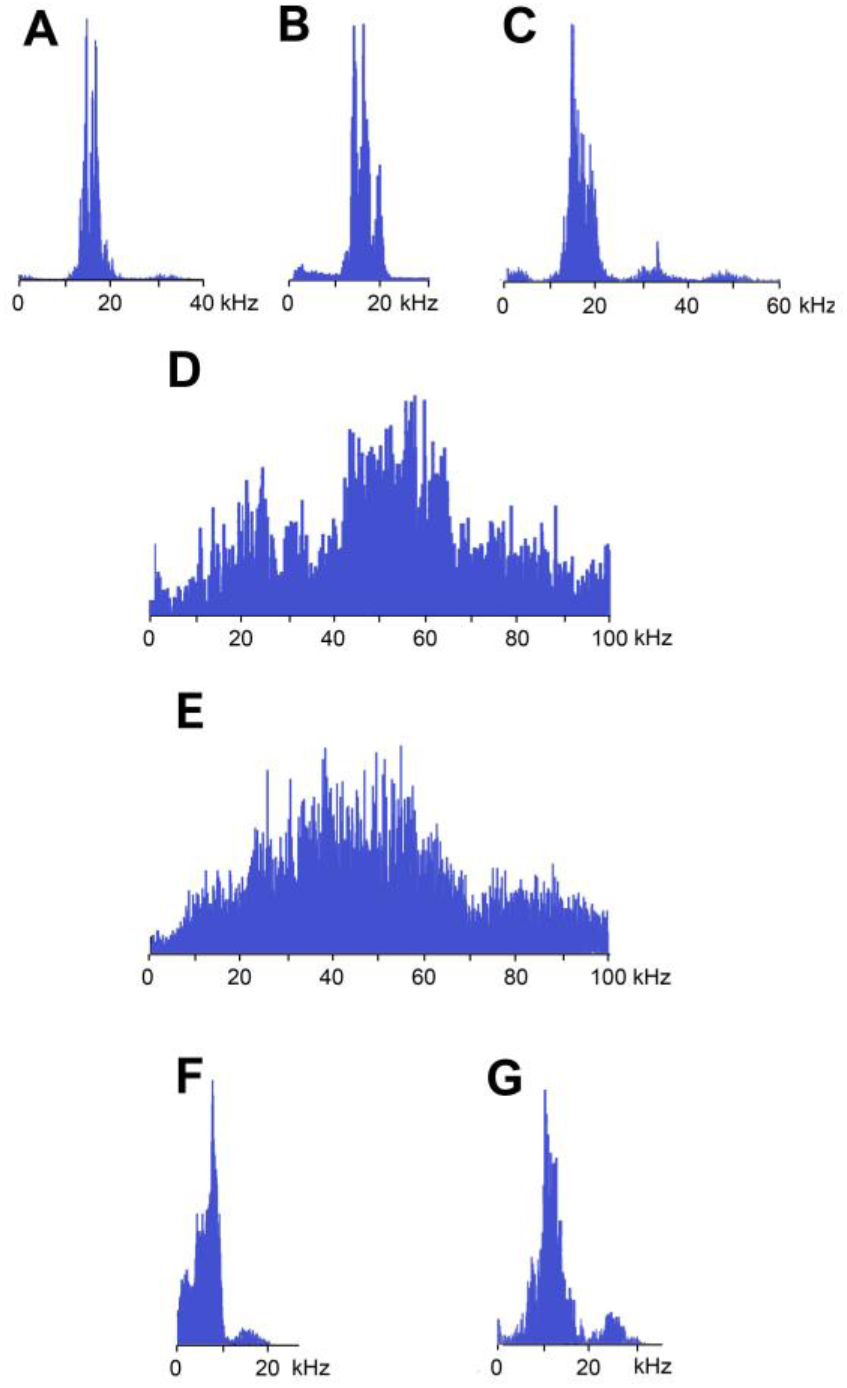
Frequency spectra (in linear scale) of sound signals of *N. nigrispina*. (A) Fast calling song. (B) Slow calling song. (C) Tegminal protest sounds. (D) Male wing protest sounds. (E) Female wing protest sounds. (F) Male mandibular protest sounds. (G) Female mandibular protest sounds

In addition to calling sounds, the acoustic repertoire of *N. nigrispina* includes three types of protest signals.

Sounds of the 1^st^ type are made by males with the help of the tegminal sound apparatus in response to mechanical stimuli, for example, to touch. As a rule, these signals are rather long series of identical single syllables (Fig. 3 E, F).Their duration is c. 40-60 ms and repetition rate is 7.7+0.8 s^−1^. However, occasionally protest signals include a series of two or even three syllables.

Frequency spectra cover the range up to 55 kHz, but the dominant frequencies are c. 14 kHz as in spectra of calling songs (Fig. 4 C).

In addition to tegminal protest sounds, males and females emit, in response to weak tactile stimuli, quiet signals with the help of their hind wings - wing protest sounds. Sometimes males and females can produce them even in the hands of a researcher. In this case, the wings spread apart and return to their original position. These movements are especially noticeable when looking at the insect from above. The elytra do not rise during sound production, which ensures their contact with the wings. Removing the elytra completely eliminates the possibility of sound producing. At the same time, wing movements are clearly visible. The SEM images (Fig. 2 D, F) show that the wings are folded in half along the midline and bear several longitudinal veins. No specialized stridulatory structures were found on the down surface of the tegmina (Fig. 2 C, E), and upper surface of the wings (Fig. 2 D, F).

Sounds of this type in males are irregular syllables of varying duration, amplitude and repetition rate. In some cases, as in Fig. 3 G, insects emit a series of syllables more or less similar in amplitude-temporal structure. Their duration is 421+44 ms, their repetition rate in a series is 1.33+0.32 s^−1^. Each male syllable begins with several clicks of increasing amplitude against the background of a low-amplitude rustling sound without amplitude modulation, followed by the same fragment without amplitude modulation (sometimes with several clicks on its background), ending with 1-2 loud short clicks (Fig. 3 I, J). In females, these signals are similar to those in males: these are either irregular syllables, or, if the insect actively reacts to the stimulus, a series of several ore or less uniform syllables with a duration of about 400 ms (Fig. 3 H, K, L). In some cases, four main parts are distinguished in their structure: the first, consisting of 5-10 clicks, the second is a part with low-amplitude high-frequency noise, lasting 100-200 ms, against the background of which individual single clicks are often recorded, and the final 3rd and 4th parts are represented by several, usually 2-4, clicks. The two final parts are separated by a pause of 50-140 ms (Fig. 4 K, L). The first part of the syllable, obviously, corresponds to the spreading of the wings, the 3rd and 4th - the final period of their return to their original position. Judging by the presence, time of appearance, and high amplitude of the initial and final clicks, the main structures of the wings (or tegmina) providing stridulation are located in their medial zone.

The frequency spectra of the wing protest sounds are also similar: they are located in both the sound and ultrasonic ranges. The dominant frequencies in males are in the band of 45-65 kHz, in females - 35-55 kHz (Fig. 4 D, E).

The third type of protest sounds that we registered in males (Fig. 3 M, N) and females (Fig. 3 O, P) during and after weak tactile stimulation are short clicks that insects produce, apparently using mandibles (Fig. 1 M, N). We observed these sounds synchronously with the movements of the mouthparts. In females, these signals consist of one or more clicks. Their duration ranges from several ms to 100 ms (on average 71.2+13.9 ms). Females produce them irregularly. Frequency spectra occupy the range of 0-30 kHz (Fig. 4 G), the dominant frequencies lie near 10 kHz.

In addition, acoustically interacting males also from time to time emit a series of low-amplitude clicks (Fig. 3 M, N), alternating with series of a calling signal or trills formed by single pulses, similar to tegminal protest signals. Having repeatedly observed these signals (they were not accompanied by drumming of any body parts on the substrate), we were able to register only one such series with the help of the equipment. The duration of clicks in it is from 2 to 17 ms (mean = 6.6 ms), their repetition rate is 3.4-4 s^−1^. Some of the clicks are double, separated by a pause of 10–20 ms. The dominant frequency in the spectrum lies in the 9-10 kHz region (Fig. 4 F) as in the spectrum of female sounds.

A video of a male and female producing protest sounds by use wings and mouthparts can be seen in Korsunovskaya, 2022 a, b.

#### Vibratory signals

When a pair is formed, up to copulation, the male and female alternately emit a series of 6-10 vibratory signals with a duration of 2.9+0.7 (male) 3.7+0.7 ms (female) (Fig. 5 C, D). The frequency of their tremulation movements repetition rate is about 4-5 s^−1^ in males and about 3 s^−1^ in females. Insects produce such signals during tremulation - vibrations of the abdomen in the vertical plane. If the range of motion is large, the male can periodically hit the substrate with the tip of the abdomen. The female usually does not touch the substrate (Korsunovskaya, 2022 c).

**Figure 5.**
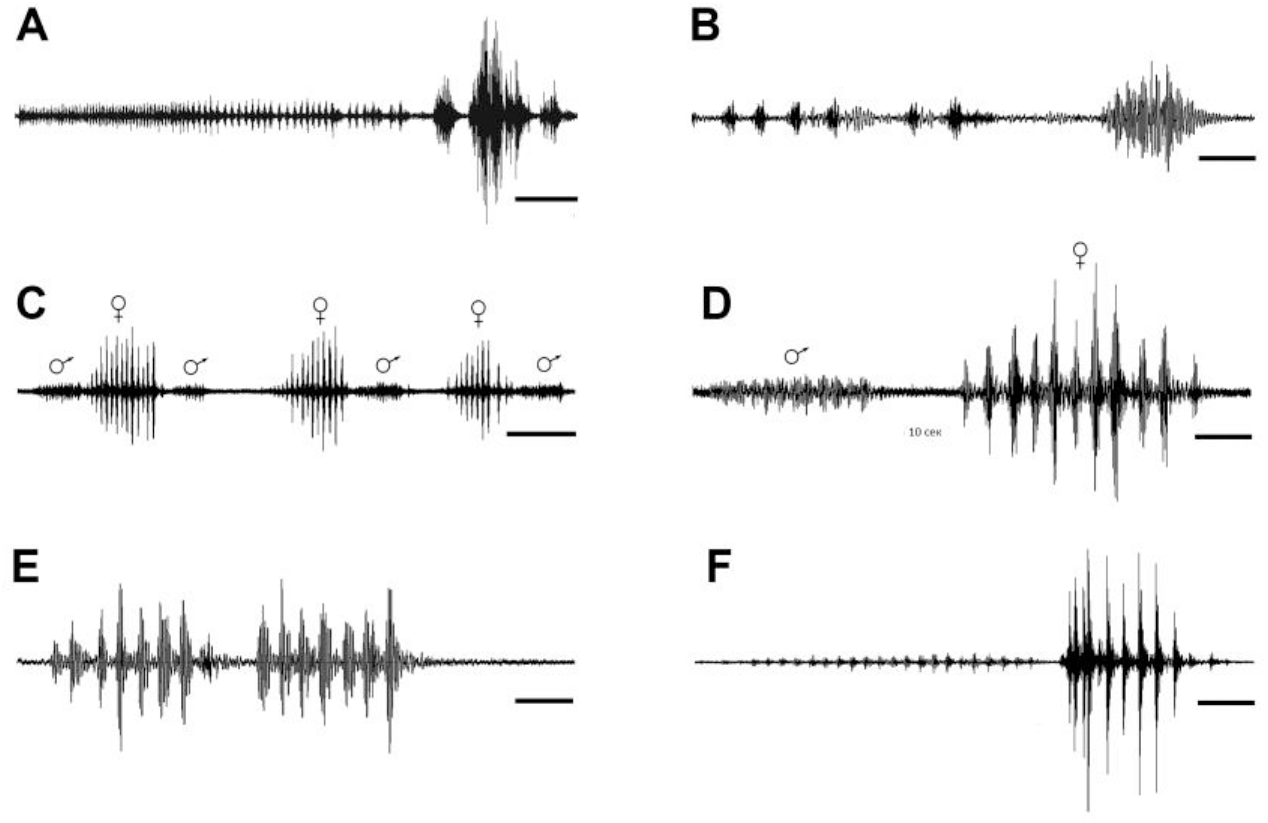
Oscillograms of the vibratory signals of *N. nigrispina*. (A, B) Vibrational component of calling song and final vibratory signal. (C, D) Male and female courtship signals. (E) Male courtship signal. (F) Two postcopulatory male signals. Time scales: 5 s (A, C) and 1 s (B, D-F)

After copulation, the female does not emit signals, and the male continues to produce rhythmic series of vibrations. They last ca. 5-7 s. From time to time, the male produces a series of oscillations of greater amplitude (Fig. 5 F), or extremely high-amplitude single blows (with duration of ca. 0.3 s) of the body against the substrate during which the insect literally jumps in place (Korsunovskaya, 2022 d).

## Discussion

The morphological characters of all currently known species of the genus *Nesoecia* are very close, and to confirm their taxonomic status it is necessary to carry out comprehensive complex studies. One of the directions of such research is the investigation of sound and vibratory signals of these species.

Comparison of the sound signals of the species studied by us with the data on the calling sounds of other representatives of *Nesoecia* (Barrientos-Lozano et al., 2020) indicates a very large similarity of their temporal and frequency characteristics. Frequency spectra of calling signals in all of the five species occupy a narrow range, and the dominant frequencies are 14-20 kHz. Unfortunately, it is difficult to compare the temporal characters of the sound signals of these katydids due to their great variability. In our opinion, the most suitable for comparison are such indicators as the shape of the syllables, the pattern of the tooth-impact frequency, as well as the syllable repetition rate in the series. Based on these parameters, both fast and slow calling signals of *N. nigrispina* are most similar to those of *N. insolita* and *N. constricta*. The main differences from *N. insolita* signals are in the shape of the first syllable, in a slow calling song and in the nature of the change in the frequency of tooth-impacts throughout the first syllable, which may indicate a differences in the structure of pars stridens and/or the speed of movement of the elytra during the first syllable in the series in both species. The sounds described by us differ from the signals of *N. constricta* by a greater sylable repetition rate in series: it is 9.1 + 0.3 s^−1^, while in *N. constricta*, judging by Fig. 123 and tab. 1 (Barrientos-Lozano et al., 2020), it is slightly more than 7 s^−1^. There are also differences in the repetition rate of the series: in *N. nigrispina* it is higher and, apparently, overlaps the variability of the signal of *N. constricta*.

The study of the vibro-acoustic signaling of *N. nigrispina* revealed several interesting facts. It turned out that these katydids during calling behavior produce not only sound, but also vibratory signals. However, unlike the neotropical *Nesonotus reticulatus* (Stumpner et al., 2013), they do this not simultaneously, but sequentially, like representatives of several genera of the tribes Cocconotini, Pleminiini (see, for example, Belwood, Morris, 1987; Montealegre, Morris, 1999). Considering that males are capable of emitting a calling signal for several hours, it cannot be concluded that the vibrational component of a calling song was the result of selection pressure from predation. Undoubtedly, it is useful for localizing a singing male, since communication takes place at night and the possibilities of visual orientation are limited.

In addition, it is possible that vibrations after a period of sound calling perform the function of an additional species-specific informative parameter. It can enhance the significance of the sound calling song as a factor in prezygotic reproductive isolation. This seems to be quite real, since data on the sound signaling of sympatric and synchronously singing katydids of the genus *Nesoecia* (Barrientos-Lozano et al., 2020), as well as a comparison of their songs with the calling signals of *N. nigrispina*, indicate a significant similarity in the temporal patterns of their songs. Another argument in favor of the importance of the vibrational component for the recognition of conspecifics is a rather high variability of calling sounds, which was noted not only in *N. nigrispina*, but also in *Xerophyllopteryx fumosa* (Brunner von Wattenwyl) (Stumpner et al., 2013).

Of particular interest is the presence of two types of audible calling signals in species of the genus *Nesoecia*: with fast and slow rhythms (Barrientos-Lozano et al., 2020, this article). As our observations have shown, insects produce fast signals during acoustic interaction. However, in some cases, these sounds can be recorded in single males for a rather long time. According to our data, katydids are most likely to emit slow signals when calling behavior. The presence of two types of calling signals in the acoustic repertoire is difficult to explain, since it is still unclear what informative elements the females use when identifying conspecific individuals. It can be assumed, by analogy with some phaneropterine bushcrickets (Spooner, 1964, 1968; Walker, 2004; Korsunovskaya, 2008; Zhantiev et al., 2017), that a fast phase with which slow calling signal of *N.nigrispina* begins or a long period of a fast signal is necessary to identify a conspecific signal and a slow phase is used during localization of its source and orientation. However, the long duration of the continuous period of slow song contradicts this assumption. Obviously, this problem can be solved only by conducting special experiments to study phonotaxis with testing various models of calling signals.

An obligatory component of courtship behavior of *N. nigrispina* is vibratory signals. This type of signal is widespread throughout the acoustically active orthopterans. It is, for example, known among the katydid subfamily Conocephalinae (see, for example, Benediktov, 2014), many Pseudophyllinae (e. g. Morris et al., 1994; Barrientos-Lozano et al., 2020), Phyllophorinae (Korsunovskaya et al., 2020). In *N. nigrispina*, these signals have a clear rhythmic pattern; they are emitted by both males and females. Apparently, along with the function of preparing for copulation and localization in the space of the sexual partner, these signals, like vibrations during calling behavior, are capable of performing the function of an additional factor in reproductive isolation. The study of courtship signals in the sympatric *Nesoecia* species may confirm or refute this hypothesis. The function of post-copulation vibratory signals appears to be to prevent the premature removal of the spermatophore by the female. Similar, extremely high-amplitude signals were described by us in the giant katydids *Siliquofera grandis* (Korsunovskaya et al., 2020).

*N. nigrispina* has a wide acoustic repertoire, including several types of signals used in agonistic relationships. We designated them as protest signals. The mechanisms for producing these sounds are different: males with a developed tegminal sound apparatus produce them in response to mechanical stimulation. Sounds of this kind, loud and prolonged, are evidently, as in katydids of the tribe Zichyini (Zhantiev et al., 1995; Elaeva, Korsunovskaya, 2012), aimed at scaring away predators. Other protest signals - wing and mandibular - are emitted by both males and females.

These are quiet signals that insects also produce in response to tactile stimuli, but males can make mandibular sounds also in response to signals from other males. In the latter case, we can, in our opinion, talk about agonistic interactions aimed at regulating intrapopulation relationships. The signals from the females are apparently also intended for conspecific individuals. Thus, the sound signaling of *N. nigrispina* is the most complex of those studied to date in representatives of Pseudophyllinae. The reasons for this development of acoustic signaling are the need for protection from predators and, apparently, the need to regulate intrapopulation relationships in various forms of competition. Judging by the absence of specialized sound apparatus used for the emission of wing and mandibular protest signals, this complication could have arisen in *N. nigrispina* during their evolution relatively recently. We assume that interactions leading to an expansion of the acoustic repertoire due to agonist signals are possible at a sufficiently high population density or in the presence of aggregations of conspecifics. However, the latter assumption requires verification in the natural habitat of this species.

## Conclusions

The known species of *Nesoecia* are morphologically very similar; therefore, the results of bioacoustic studies are very important for confirming the species status of the congenerics. The study of the vibro-acoustic signaling of *N. nigrispina* showed that these katydids, during calling and courtship, produce not only sound, but also vibratory signals. The repertoire of sound signals of *N. nigrispina* is the most extensive among the representatives of the family Pseudophyllidae. It includes calling song of two types and 3 types of protest signals. Males, which, like other katydids, have tegminal stridulatory organ, produce sounds of all types. However, the emission of protest signals of the 2nd and 3rd types is carried out with the help of the wings and mouth organs (apparently, mandibles). Females can also produce Type 2 and Type 3 protest sounds in the same way as males. No specialized structures for the emission of wing and mandibular sounds have been identified, which may indicate that the expansion of the acoustic repertoire in the evolution occurred relatively recently. Perhaps this arose in connection with the need not only to perform a protective function against the attack of predators, but also to regulate the agonistic intrapopulation relations. Vibratory signals are emitted by individuals of both sexes during courtship and males, completing the calling signal cycle and after copulation. It is possible that vibratory signals are an additional factor in reproductive isolation in sympatric species, since the calling sound signals in representatives of the genus *Nesoecia* are similar and exhibit significant variability. What type and what parameters of the calling signal the female uses when identifying a conspecific sexual partner should be revealed by special studies.

## Acknowledgements

We are grateful to M. Berezin and T. Kompantseva for providing insects for katydid culture, S. Farisenkov for providing us with equipment for high-speed video recording and infrared lighting, A. Gorokhov for help in species identification of *Nesoecia*. This study was supported by the Russian Foundation for Basic Research, project 19-04-00104.

## Notes

### Competing Interest Statement

The authors have declared no competing interest.

## References

Barrientos-Lozano L., Rocha-Sanchez A.Y., Fernández-Azuara G. de J., Sánchez-Reyes U. J., Almaguer-Sierra P. 2020. Cryptic new species of *Nesoecia* Scudder, 1893 (Orthoptera: Tettigoniidae; Pseudophyllinae) from northeastern, Mexico. Zootaxa. 4859(4):451–486 DOI: 10.11646/zootaxa.4859.4.1

Belwood J. J. 1990. Anti-predator defences and ecology of neotropical katydids, especially the Pseudophyllinae. In: Bailey W. J., Rentz D. C. F., Editors. The Tettigoniidae: Biology, Systematics and Evolution. pp. 8–26. Springer-Verlag.

Belwood J. J., Morris G. K. 1987. Bat predation and its influence on calling behavior in Neotropical katydids. Science, Wash. 238: 64–67. DOI: 10.1126/science.238.4823.64

Benediktov A. A. 2014. Vibro-acoustical signals of the meadow katydids from the subfamily Conocephalinae (Orthoptera, Tettigoniidae) in the European part of Russia. Moscow University Biological Sciences Bulletin. 69(4):180–183 DOI https://doi.org/10.3103/S0096392514040026

Cigliano, M. M., Braun, H., Eades, D. C. & Otte, D. (2021) Orthoptera Species File. Version 5.0/5.0. Available from: http://Orthoptera.SpeciesFile.org/ (accessed 30 December 2021)

Elaeva N. F., Korsunovskaya O. S. 2012. Environmental features and sound signaling of relict katydid Deracantha onos (Pallas, 1772) (Orthoptera, Tettigoniidae, Bradyporinae). Moscow University Biological Sciences Bulletin. 67 (3):105–111 DOI https://doi.org/10.3103/S0096392512020046

Heller K.-G. 1996. Unusual abdomino-alary, defensive stridulatory mechanism in the bushcricket Pantecphylus cerambycinus (Orthoptera: Tettigoniidae, Pseudophyllidae). Journal of Morphology. 227(1):81–86 DOI: 10.1002/(SICI)1097-4687(199601)227:1%3C81::AID-JMOR%63E3.0.CO;2-S

Heller K.-G., Baker E., Ingrisch S., Korsunovskaya O. S. l, Liu C. X., Riede K., Warchałowska-Šliwa E. 2021. Bioacoustics and systematics of *Mecopoda* (and related forms) from South East Asia and adjacent areas (Orthoptera, Tettigonioidea, Mecopodinae) including some chromosome data. Zootaxa, 5005(2):101–144. DOI: https://doi.org/10.11646/zootaxa.5005.2.1

Korsunovskaya O. S. 2008. Acoustic signals in katydids (Orthoptera, Tettigoniidae). Communication I. Entomological Review, 88(9):1032–1050. DOI: https://doi.org/10.1134/S0013873808090029

Korsunovskaya O. 2022 a. Video of female wing and mandibular protest sounds of the katydid, *Nesoecia nigrispina* (Orthoptera, Pseudophyllinae). Zenodo. DOI: https://doi.org/10.5281/zenodo.5913684

Korsunovskaya O. 2022 b. Video of male wing and mandibular protest sounds of the katydid, *Nesoecia nigrispina* (Orthoptera, Pseudophyllinae). Zenodo. DOI: https://doi.org/10.5281/zenodo.5913615

Korsunovskaya O. 2022 c. Video of precopulatory tremulation during courtship in *Nesoecia nigrispina* (Orthoptera, Pseudophyllinae). Zenodo. DOI: https://doi.org/10.5281/zenodo.5913721

Korsunovskaya O. 2022 d. Video of male postcopulatory tremulation of the katydid, *Nesoecia nigrispina* (Orthoptera, Pseudophyllinae). Zenodo. DOI: https://doi.org/10.5281/zenodo.5913738

Korsunovskaya O., Berezin M., Heller K.-G., Tkacheva E., Kompantseva T., Zhantiev R. 2020. Biology, sounds and vibratory signals of hooded katydids (Orthoptera: Tettigoniidae: Phyllophorinae). Zootaxa. 4852(3): 309–322. DOI: https://doi.org/10.11646/zootaxa.4852.3.3

Montealegre-Z. F, Guerra P. A., Morris G. K. 2009. *Panoploscelis specularis* (Orthoptera: Tettigoniidae: Pseudophyllinae): extraordinary female sound generator, male description, male protest and calling signals. Journal of Orthoptera Research 2003,12(2): 173–181. DOI: 10.1665/1082-6467(2003)012[0173:PSOTPE]2.0.CO;2

Montealegre-Z F., Morris G. K. 1999. Songs and Systematics of some Tettigoniidae from Colombia and Ecuador, part I. Pseudophyllinae (Orthoptera). Journal of Orthoptera Research. 8(8):163–236. DOI:10.2307/3503439

Morris G. K., Mason A. C., Wall P., Belwood J. J. 1994. High ultrasonic and tremulation signals in neotropical katydids (Orthoptera: Tettigoniidae). Journal of Zoology. Lond. 233(1): 129–163. DOI:10.1111/j.1469-7998.1994.tb05266.x

Rajaraman K., Godthi V., Pratap R., Balakrishnan R. 2015. A novel acoustic-vibratory multimodal duet. Journal of Experimental Biology. 218: 3042–3050. DOI: https://doi.org/:10.1242/jeb.122911

Roemer H., Lang A., Hartbauer M. 2010. The signaller’s dilemma: a cost–benefit analysis of public and private communication. PLoS ONE 5(10): e13325. DOI: https://doi.org/:10.1371/journal.pone.0013325

Spooner J. D. 1964. The texas bush crickets - its sounds and their significance // Animal Behaviour. 12(2-3): 235–244. DOI:10.1016/0003-3472(64)90007-7

Spooner J. D. 1968. Pair-forming acoustic systems of phaneropterine katydids (Orthoptera, Tettigoniidae) // Animal Behaviour. 16(2-3): 197–212. DOI: 10.1016/0003-3472(68)90001-8

Stumpner A., Dann A., Schink M., Gubert S., Hugel S. 2013. True katydids from Guadeloupe: Acoustic signals and functional considerations of song production. Journal of Insect Science 13(1):157. DOI:10.1673/031.013.15701.

Walker T. J. 2004. The *uhleri* group of the genus *Amblycorypha* (Orthoptera: Tettigoniidae): extraordinarily complex songs and new species // Journal of Orthoptera Research. 13 (2): 169–183. DOI:10.1665/1082-6467(2004)013[0169:TUGOTG]2.0.CO;2

Zhantiev R., Korsunovskaya O., Benediktov A. 2017. *Acoustic signals of the bush-crickets Isop hya Br.-W. 1878 (Orthoptera: Phaneropteridae) from Eastern Europe, Caucasus and* adjacent territories. European Journal of Entomology, 114: 301–311. DOI: 10.14411/eje.2017.037

Zhantiev, R. D., Korsunovskaya, O. S., and Byzov, S. D., 1995. Acoustic communication of arid katydids (Bradyporidae, Deracanthinae). [In Russian]. Zooogicheskii Zhurnal 74(9):58–71.

